# Isoform switching as a mechanism of acquired resistance to isocitrate dehydrogenase inhibition

**DOI:** 10.1101/381954

**Authors:** James J. Harding, Maeve A. Lowery, Alan H. Shih, Juan M. Schvartzman, Shengqi Hou, Christopher Famulare, Minal Patel, Mikhail Roshal, Richard K. G. Do, Ahmet Zehir, Daoqi You, S. Duygu Selcuklu, Agnes Viale, Martin S. Tallman, David M. Hyman, Ed Reznik, Lydia W.S. Finley, Elli Papaemmanuil, Alessandra Tosolini, Mark G. Frattini, Kyle J. MacBeth, Guowen Liu, Bin Fan, Sung Choe, Bin Wu, Yelena Y. Janjigian, Ingo K. Mellinghoff, Luis A. Diaz, Ross L. Levine, Ghassan K. Abou-Alfa, Eytan M. Stein, Andrew M. Intlekofer

## Abstract

Somatic mutations in cytosolic or mitochondrial isoforms of isocitrate dehydrogenase (IDH1 or IDH2, respectively) contribute to oncogenesis via production of the metabolite 2-hydroxyglutarate (2HG). Isoform-selective IDH inhibitors suppress 2HG production and induce clinical responses in patients with IDH1- and IDH2-mutant malignancies. Despite the promising activity of IDH inhibitors, the mechanisms that mediate resistance to IDH inhibition are poorly understood. Here, we describe four clinical cases that identify mutant IDH isoform switching, either from mutant IDH1 to mutant IDH2 or vice versa, as a mechanism of acquired clinical resistance to IDH inhibition in solid and liquid tumors.

**Significance:** IDH-mutant cancers can develop resistance to isoform-selective IDH inhibition by “isoform switching” from mutant IDH1 to mutant IDH2 or vice versa, thereby restoring 2-hydroxyglutarate (2HG) production by the tumor. These findings underscore a role for continued 2HG production in tumor progression and suggest therapeutic strategies to prevent or overcome resistance.

## Introduction

Somatic mutations in isocitrate dehydrogenase (IDH) enzymes are observed in a wide spectrum of human cancers, most commonly gliomas^1,2^, myeloid malignancies^3^, chondrosarcomas^4^, and intrahepatic cholangiocarcinomas (ICC)^5^. IDH enzymes normally function as components of the tricarboxylic acid cycle, catalyzing the interconversion of isocitrate and alpha-ketoglutarate (αKG)^6^. Cancer-associated IDH mutations occur at catalytic-site arginine residues in cytoplasmic IDH1 (R132) and mitochondrial IDH2 (R140 and R172), altering the enzymatic activity to promote efficient reduction of αKG to the ‘oncometabolite’ 2-hydroxyglutarate (2HG)^7,8^. Cellular accumulation of 2HG inhibits αKG-dependent dioxygenases that mediate histone and DNA demethylation, leading to a repressive chromatin landscape that disrupts normal cellular differentiation and contributes to oncogenesis^9–12^.

Importantly, the effects of 2HG on chromatin and cell differentiation are largely reversible^11^, therefore drugs that inhibit 2HG production by mutant IDH represent potential therapies for IDH-mutant malignancies^13^. Small-molecule inhibitors of IDH1 or IDH2 inhibit 2HG production by IDH-mutant enzymes, induce cellular differentiation, and inhibit cancer growth in preclinical models^6,13^. In phase I/II trials, the mutant IDH1 inhibitor ivosidenib (AG-120) and the mutant IDH2 inhibitor enasidenib (AG-221) induced clinical responses in patients with relapsed/refractory IDH-mutant myeloid malignancies^14,15^. Clinical trials with ivosidenib are underway for the treatment of relapsed/refractory IDH1-mutant solid tumors, including glioma, ICC, and chondrosarcoma^16^.

The mechanisms that mediate response and resistance to small-molecule IDH inhibition remain poorly understood^17–19^. Here, we identify mutant IDH isoform switching, either from cytoplasmic mutant IDH1 to mitochondrial mutant IDH2 or vice versa, as a mechanism of acquired clinical resistance to IDH inhibition. The first two cases describe patients with relapsed/refractory IDH1 R132C-mutant AML who achieved durable remissions in response to the mutant IDH1 inhibitor ivosidenib followed by leukemic progression on therapy, rise of blood 2HG levels, and emergence of new IDH2 R140Q mutations. The third case describes a patient with treatment-refractory IDH1 R132C-mutant intrahepatic cholangiocarcinoma who attained a sustained partial response to ivosidenib followed by progression of disease associated with acquisition of a new IDH2 R172V mutation. The fourth case describes a patient with relapsed/refractory IDH2 R140Q-mutant acute myeloid leukemia who achieved a durable remission to the mutant IDH2 inhibitor enasidenib, followed by progression on therapy associated with emergence of a new IDH1 R132C mutation that was sensitive to combined IDH1/2 blockade with AG-881. These findings provide evidence of selective pressure to maintain 2HG production in IDH-mutant malignancies, as well as suggest potential strategies for disease monitoring and therapies that might overcome acquired resistance to IDH inhibition.

## Results

### Case Reports 1 and 2

A 54-year-old man with normal karyotype AML relapsed 100 days after allogeneic bone marrow transplantation with progressive pancytopenia and a bone marrow biopsy showing 30% leukemic blasts (Figures 1A and B). Targeted NGS of bone marrow cells using a microdroplet-PCR assay^20^ demonstrated the presence of an IDH1 R132C mutation and two DNMT3A mutations (Figure 1C and Supplementary Tables 1 and 4). The patient began treatment with ivosidenib 500 mg orally daily, with a complete remission evident after three 28-day cycles of therapy. After completing twelve 28-day cycles of ivosidenib, the blast count and blood 2HG levels began to rise and a new IDH2 R140Q mutation was detected (Figures 1A, C, D). Ivosidenib was discontinued and enasidenib was started. After several days of treatment with enasidenib, the patient developed fevers and hypoxia suspected to be secondary to IDH inhibitor differentiation syndrome^21^; enasidenib was discontinued.

**Figure 1.**
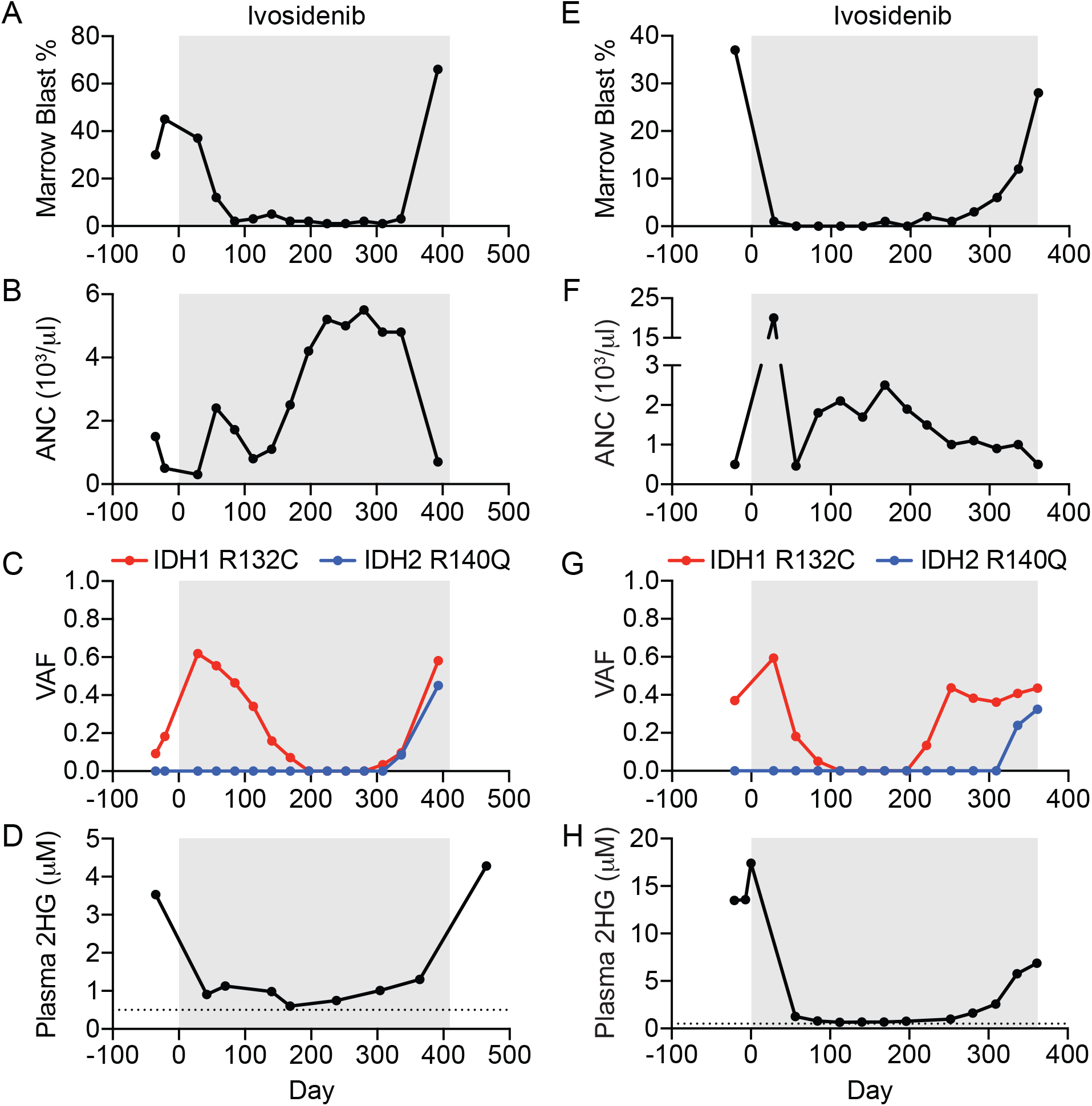
Acquired resistance to mutant IDH1 inhibition associated with emergence of oncogenic IDH2 mutations in acute myeloid leukemia. Clinical and laboratory features of two patients (Case 1 = A–D; Case 2 = E–H) with IDH1 R132C-mutant acute myeloid leukemia treated with the mutant IDH1 inhibitor ivosidenib (gray boxes), including (A, E) bone marrow blast percentage, (B, F) absolute neutrophil count (ANC), (C, G) variant allele frequency (VAF) for IDH1 and IDH2 mutations identified by targeted next-generation sequencing of bone marrow cells, and (D, H) plasma 2-hydroxyglutarate (2HG) concentration measured by gas chromatography–mass spectrometry.

A 72-year-old man presented with AML arising from pre-existing JAK2 V617F-mutant myelofibrosis. For several years prior, the myelofibrosis had been treated successfully with single-agent ruxolitinib, then combination therapy with ruxolitinib plus decitabine. However, at the time of presentation with secondary AML, there were 37% blasts in the bone marrow and the patient was neutropenic (Figures 1E and F). Targeted NGS of bone marrow mononuclear cells using a microdroplet-PCR assay^20^ demonstrated the presence of JAK2 V617F and IDH1 R132C mutations (Figure 1G), as well as mutations in ASXL1, EZH2, and RUNX1 (Supplementary Tables 1 and 5). The patient began treatment with ivosidenib 500 mg orally daily, and a complete response was evident after one 28-day cycle of therapy. The IDH1 mutation became undetectable after four 28-day cycles of ivosidenib, but then reappeared after the eighth 28-day cycle (Figure 1G). The patient remained in complete morphologic remission until the start of the twelfth cycle when the bone marrow blasts increased to 12% then 28% four weeks later (Figure 1E). The increase in AML blasts was associated with the emergence of a new IDH2 R140Q mutation and a rise in the serum 2HG levels (Figures 1G and H). Ivosidenib was discontinued and the patient subsequently pursued treatment with low-dose cytarabine and venetoclax^22^.

### Case Report 3

A 79-year-old woman with AJCC Stage IV (T3N1M1) intrahepatic cholangiocarcinoma (ICC) presented for evaluation. One month prior to presentation she had developed anorexia, unintentional weight loss, and abdominal distention. Cross-sectional imaging with computed tomography revealed an 8×5×7.5 cm hypoattenuating mass with peripheral enhancement and capsular retraction in the right hepatic lobe, multiple hepatic satellite tumors, and extensive retroperitoneal lymphadenopathy. Core biopsy of the dominant right hepatic mass revealed a poorly differentiated adenocarcinoma with immunohistochemical markers consistent with a primary biliary tumor. An analysis of genomic DNA from the biopsy specimen using targeted NGS revealed an IDH1 R132C mutation and no other detectable mutations (Supplemental Tables 2 and 6).

After documented radiographic progression on multiple standard and investigational agents, the patient began treatment with ivosidenib 500 mg orally daily. Computed tomographic scans prior to treatment and then serially over the course of treatment demonstrated a partial response to treatment with a 50% decrease in disease burden (Figures 2A and B). An escape lesion was identified at day 392, and this lesion continued to enlarge on a follow-up scan at day 446, despite a further decrease in size of the original tumor (Figure 2B). Ivosidenib was discontinued and a post-progression biopsy of the enlarging lesion was performed. Sequencing of the resistant tumor confirmed the presence of the initial IDH1 R132C in addition to a new IDH2 R172V mutation, CDKN2A/B loss, and several other alterations of uncertain significance (Figure 2C and Supplemental Tables 3 and 6). Subsequently, the patient was treated with pembrolizumab, a monoclonal antibody to program death receptor-1, with slow progression of disease on serial imaging and clinical progression with worsening ascites and diminution of ECOG performance status. She then began treatment with the mutant IDH2 inhibitor enasidenib but within 8 weeks underwent clinical deterioration and expired.

**Figure 2.**
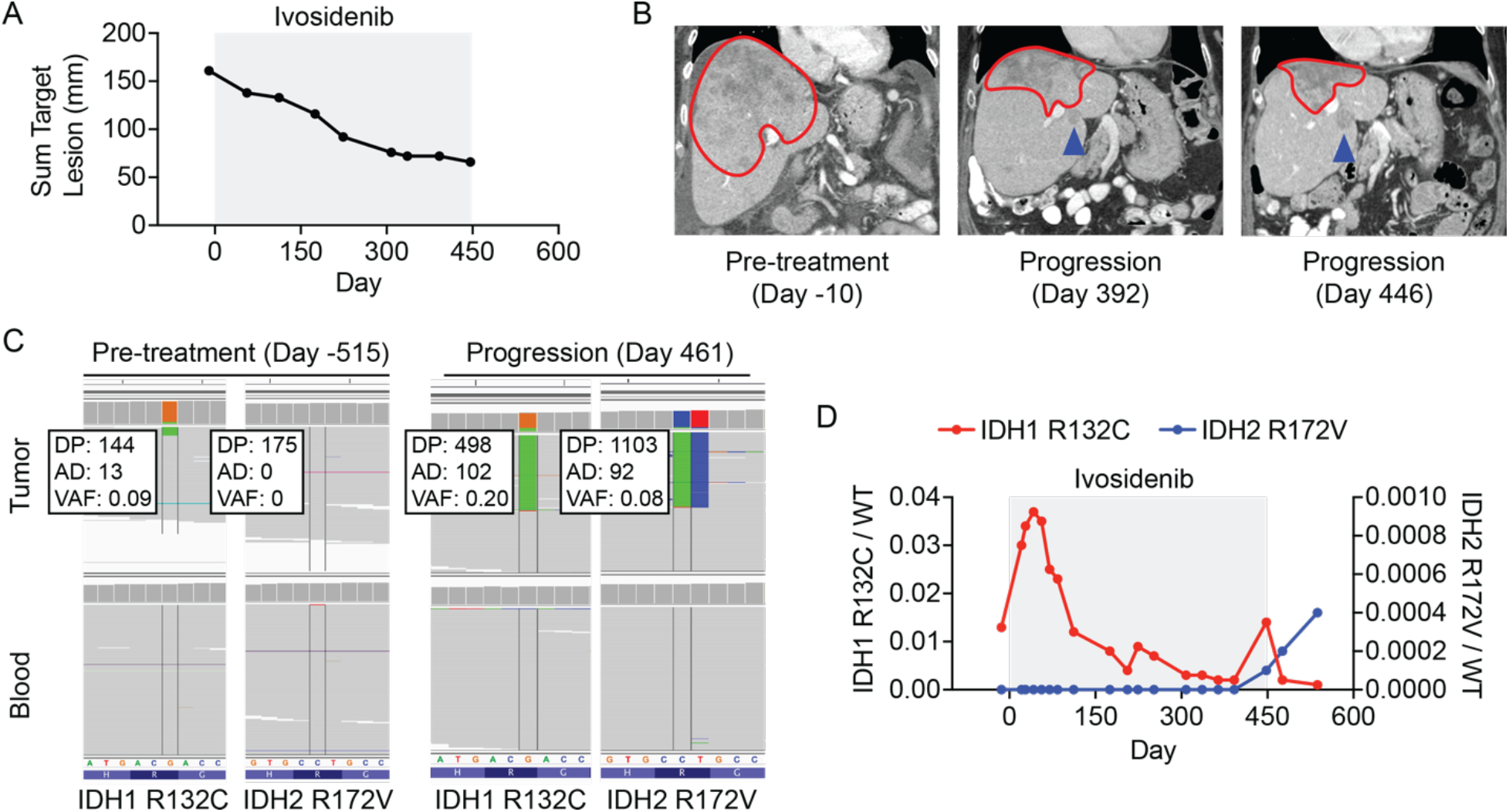
Acquired resistance to mutant IDH1 inhibition associated with emergence of an oncogenic IDH2 mutation in cholangiocarcinoma. Radiographic and laboratory features of a patient (Case 3) with IDH1 R132C-mutant intrahepatic cholangiocarcinoma treated with the mutant IDH1 inhibitor ivosidenib (gray box), including (A) relative change in cumulative tumor size over time, (B) serial cross-sectional imaging of the tumor mass in the liver, (C) Integrative Genomics Viewer (IGV) images highlighting relevant regions of IDH1 and IDH2 in pretreatment (Day -515) and progression biopsy (Day 461) sequenced by MSK-IMPACT, and (D) ratios of sequencing reads for IDH1 R132C/WT (red, left axis) or IDH2 R172V/WT (blue, right axis) as assessed by droplet digital PCR of cell-free DNA isolated from peripheral blood. For (B), the red line demarks the primary tumor and the blue arrow indicates the escape lesion. DP, total depth. AD, depth of alternative allele. VAF, variant allele frequency.

Serial monitoring of cell-free DNA by droplet digital PCR (ddPCR) for IDH1 R132C and IDH2 R172V demonstrated the presence of the IDH1 R132C mutant allele prior to ivosidenib, with an allelic fraction that decreased over the course of ivosidenib therapy (Figure 2D). In contrast, the IDH2 R172V mutation first became detectable at day 448 coinciding with radiographic disease progression (Figure 2D). Measurement of 2HG levels by liquid chromatography–mass spectrometry (LC-MS) demonstrated increased 2HG within the escape tumor lesion (Figure S1A). However, possibly due to the small size of this lesion, there was no detectable increase in plasma 2HG at the time of progression (Figure S1B). Introduction of the IDH2 R172V mutation into IDH1 R132C-mutant chondrosarcoma cells restored 2HG production that was resistant to ivosidenib, but susceptible to combined treatment with ivosidenib and enasidenib (Figure S1C).

### Case Report 4

A previously healthy 70-year-old man was found to be pancytopenic on routine bloodwork. Bone marrow aspiration and core biopsy demonstrated a hypercellular marrow with dysplastic neutrophils, erythroid precursors, and megakaryocytes. Myeloblasts were increased and comprised 15% of the marrow cellularity. These morphologic findings were consistent with myelodysplastic syndrome (MDS), subtype refractory anemia with excess blasts 2 (RAEB-2). Cytogenetics and fluorescence in situ hybridization (FISH) were normal. Clinical amplicon-based genotyping of bone marrow cells demonstrated the presence of an IDH2 R140Q mutation without mutation of NPM1, CEBPA, FLT3, or KIT (not shown). After 4 cycles of azacytidine, pancytopenia persisted (Figures S2A-C). Repeat bone marrow evaluation demonstrated a hypercellular marrow with persistent trilineage dysplasia (Figure S2D) and increased myeloid blasts comprising up to 20% of the marrow cellularity (Figure 3A). Targeted next-generation sequencing (NGS) of bone marrow cells using a microdroplet-PCR assay^20^ confirmed persistence of the IDH2 R140Q mutation (Figure 3B and Supplemental Tables 1 and 7). Based on these findings, the patient met the diagnostic criteria for IDH2-mutant secondary acute myeloid leukemia (AML) evolved from MDS.

**Figure 3.**
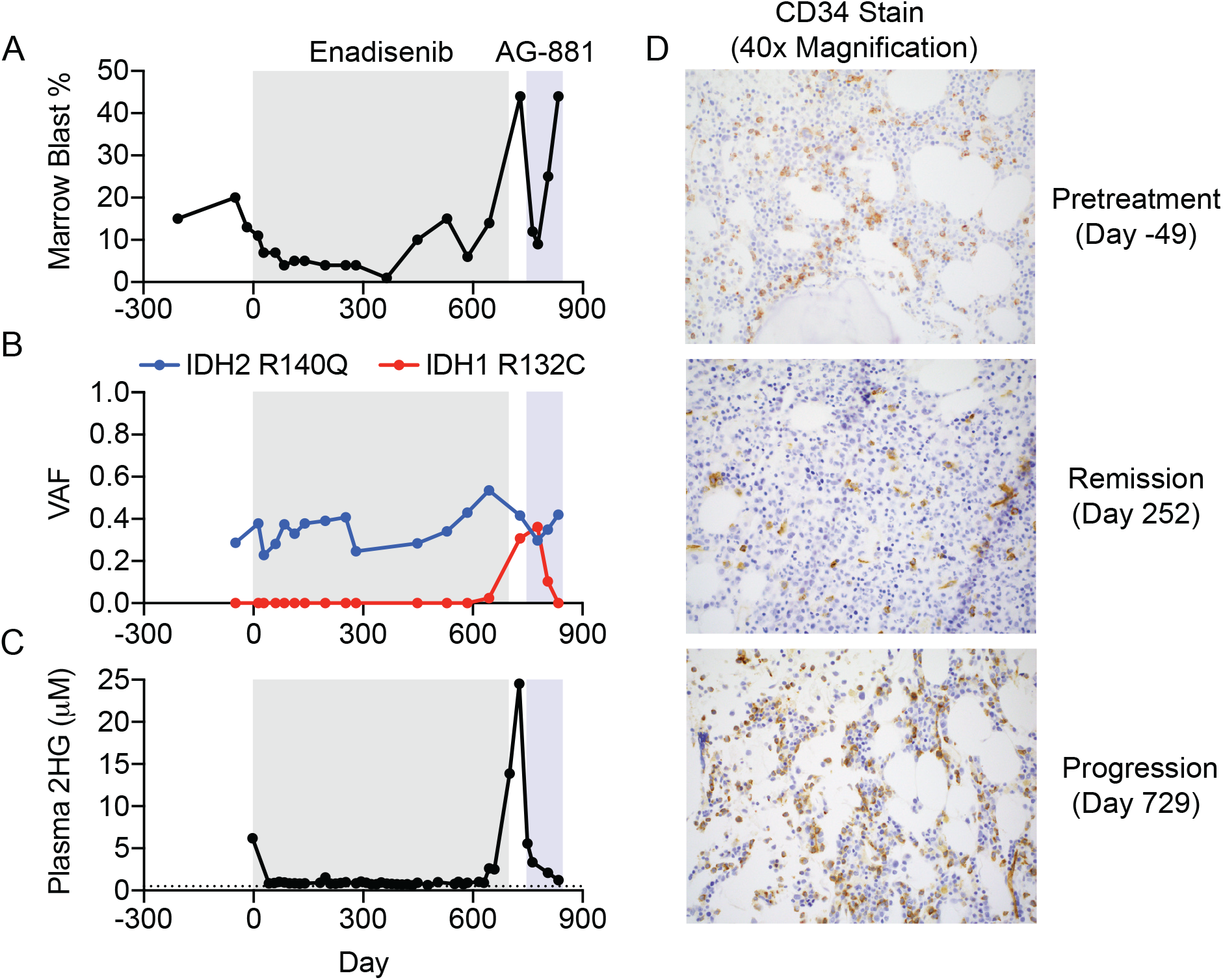
Acquired resistance to mutant IDH2 inhibition associated with emergence of an oncogenic IDH1 mutation in acute myeloid leukemia. Clinical, laboratory, and pathologic features of a patient (Case 4) with IDH2 R140Q-mutant acute myeloid leukemia treated with the mutant IDH2 inhibitor enasidenib (gray box) and the dual IDH1/2 inhibitor AG-881 (blue box), including (A) bone marrow blast percentage, (B) variant allele frequency (VAF) for IDH1 and IDH2 mutations identified by targeted next-generation sequencing of bone marrow cells, (C) plasma 2-hydroxyglutarate (2HG) concentration measured by gas chromatography–mass spectrometry (Day -3 by liquid chromatography–mass spectrometry), and (D) CD34 immunohistochemical staining (identifies leukemic blasts) of bone marrow cells at indicated points in the disease course.

The patient began treatment with the mutant IDH2 inhibitor enasidenib 100 mg orally twice daily. Monthly bone marrow evaluation demonstrated a progressive decrease in the myeloid blast percentage with a complete response (blasts <5%) evident after completing three 28-day cycles of therapy (Figures 3A and S2D). Although the pancytopenia persisted, there was a modest improvement in the absolute neutrophil count (Figures S2A-C). The variant allele frequency (VAF) for IDH2 R140Q remained in the ~20-40% range (Figure 3B), consistent with the observation that enasidenib promotes differentiation of IDH2-mutant blasts rather than cytotoxicity^14,17^. Serial measurements of plasma 2HG concentration demonstrated a decrease from the pre-treatment level of 6.2 μM to levels ranging from 0.5-1.5 μM while on therapy with enasidenib (Figure 3C).

The patient remained in complete remission until the start of cycle 17 of enasidenib (day 448), at which time the bone marrow blast percentage increased to 10% (Figure 3A), leukemic blasts appeared in the blood (not shown), and two CEBPA mutations became detectable by NGS of bone marrow cells (Supplementary Table 7). On day 497, the dose of enasidenib was increased to 200 mg orally twice daily, resulting in a transient decrease in bone marrow blasts (Figure 3A). However, eight weeks later there was an increase in bone marrow blasts accompanied by a progressive rise in plasma 2HG concentration (Figures 3A, C, and D). Targeted NGS of bone marrow cells demonstrated the appearance of a new IDH1 R132C mutation (Figure 3B) coinciding with the leukemic progression and rise in plasma 2HG levels.

Enasidenib was discontinued on day 700, and the patient began therapy on a clinical study of the dual IDH1/2 inhibitor AG-881 at 400 mg orally daily. There were rapid decreases in the plasma 2HG concentration, bone marrow blast percentage, and variant allele fraction (VAF) for IDH1 R132C (Figures 3A-C). These effects were transient, as the patient underwent a prolonged hospitalization for *Clostridium difficile* infection and gastrointestinal bleeding and ultimately expired approximately 3 months after starting therapy with AG-881. In the interval prior to expiration, there was progression of leukemic blasts despite ongoing suppression of blood 2HG levels.

## Discussion

We have identified mutant IDH isoform switching as a mechanism of acquired resistance to IDH-targeted therapy. Both IDH1 and IDH2 mutations are observed in liquid tumors, whereas in solid tumors IDH1 mutations vastly outnumber IDH2 mutations^6^. The biological underpinning for the preferential selection of cytosolic IDH1 mutation versus mitochondrial IDH2 mutation in different cancer types remains poorly understood. Experimental evidence suggests that the level of 2HG production can vary based on the subcellular location of the IDH mutation^23^. Our findings suggest that the selective pressure of inhibiting mutant IDH activity in one subcellular compartment might provide an advantage for malignant subclones that acquire unimpeded mutant IDH activity in another subcellular compartment. It is unclear whether the resistance alleles were acquired during the course of treatment or were present prior to treatment in a rare subclone that was below the limit of detection of the NGS and ddPCR assays. Although the variant allele frequencies of the IDH mutations raise the possibility that the IDH1 and IDH2 mutations might be acquired within the same subclone, single cell analysis will be required to delineate whether resistance is due to serial or parallel mutation acquisition. Likewise, it will be important to determine whether additional co-occurring genomic alterations might contribute to acquired resistance to IDH inhibition.

Mutant IDH isoform switching represents a mechanism of acquired resistance to isoform-selective IDH targeted therapy with therapeutic import. Experimental evidence demonstrated that a cancer cell line expressing both mutant IDH1 and mutant IDH2 exhibited 2HG production that was resistant to monotherapy, yet sensitive to combined inhibition of both isoforms (Fig S1C). The patient in Case Report 4 exhibited potent inhibition of 2HG production and a transient clinical response to the dual IDH1/2 inhibitor AG-881, although there was subsequent 2HG-independent disease progression resulting from unknown molecular mechanisms. Alternatively, a strategy of co-targeting of mutant IDH1 and IDH2 from the outset might delay or prevent resistance and should be explored in clinical trials. As larger numbers of patients with IDH-mutant malignancies are treated with isoform selective IDH inhibitors, it will be important to establish the frequency of isoform switching and to elucidate other mechanisms of de novo and acquired resistance^17–19^. These cases underscore the potential utility of serial NGS and 2HG measurement in patients treated with IDH inhibitors as means to monitor response, define resistance mechanisms, and inform treatment recommendations.

## Acknowledgements

We thank members of the Finley, Intlekofer, Levine, Papaemmanuil, and Reznik laboratories for helpful discussions. A.M.I. is supported by the NIH/NCI (K08 CA201483), Burroughs Wellcome Fund (CAMS 1015584), Damon Runyon Cancer Research Foundation (CI 95-18), Leukemia & Lymphoma Society (Special Fellow 3356-16), Susan & Peter Solomon Divisional Genomics Program, Steven A. Greenberg Fund, and Cycle for Survival. A.H.S is supported by the NIH/NCI (K08 CA181507) and Leukemia & Lymphoma Society. J.M.S. is a Hope Funds for Cancer Research Postdoctoral Fellow. The work was also supported, in part, by the Leukemia & Lymphoma Society Specialized Center of Research Program (7011-16; A.M.I., C.B.T.), a Translational and Integrative Medicine Research Fund (TIMRF) grant (A.H.S., E.M.S.), and grants from the NIH, including R35 CA197594-01A1 (R.L.L.), U54 OD020355 (R.L.L.), and the Memorial Sloan Kettering Cancer Center Support Grant (NIH P30 CA008748). We acknowledge the use of the Integrated Genomics Operation Core, funded by the Memorial Sloan Kettering Cancer Center Support Grant (NIH P30 CA008748), Cycle for Survival, and the Marie-Josée and Henry R. Kravis Center for Molecular Oncology.

## Author Contributions

J.J.H., M.A.L., E.M.S. and A.M.I. conceived the project and analyzed the data. J.J.H., E.M.S. and A.M.I. wrote the manuscript. J.M.S., S.H., C.F., M.P., M.R., R.K.G.D., A.Z., D.Y., S.D.S., A.V., M.T., D.M.H., E.R., L.W.S.F., E.P., A.T., M.G.F., K.M., G.L., B.F., S.C., B.W., Y.Y.J., I.K.M., L.A.D., R.L.L. and G.K.A. assisted with collection and management of clinical data and pathologic/molecular assessment of biospecimens. All authors read and approved the manuscript.

## Methods

### Clinical specimens

The patients described were enrolled on the phase I/II studies NCT01915498^14^, NCT02073994, or NCT02074839. Enrollment was open to all patients with relapsed or refractory acute myeloid leukemia with a mutation in IDH1 (NCT02074839) or IDH2 (NCT01915498) or patients with advanced solid tumors with a mutation in IDH1 (NCT02073994). IDH mutations were identified locally and confirmed centrally. Patients were required to be 18 years or older at the time of study entry. Both men and women were enrolled on the studies. Patients were required to have acceptable performance status and adequate organ function as defined in the study protocols. Clinical data, blood, bone marrow, or tumor biopsy specimens were obtained after receiving written informed consent from patients. Approval was obtained from the Institutional Review Board at each institution participating in these clinical trials. Additional consent was obtained from participants at Memorial Sloan Kettering Cancer Center with analyses performed on the institutional biobanking protocol approved by the Institutional Review Board. Patient biospecimens were anonymized by creating unique identifiers with no associated PHI and keeping the key on a password-protected server. Data collection and research was performed in compliance with all relevant ethical regulations for human research participants. Absolute neutrophil count, hemoglobin concentration, platelet count, and blast percentage were determined by standard clinical assays. Targeted next-generation sequencing and determination of variant allele frequencies (VAF) were performed by Raindance™ microdroplet PCR^20^ for AML or MSK-IMPACT™ hybrid capture^24^ for cholangiocarcinoma as previously described (see Supplementary Tables 1-3 for gene lists).

### Cell-free DNA (cfDNA) isolation and sequencing

Whole blood was collected in 10-ml Cell-Free DNA BCT tubes (STRECK) and centrifuged at 800 g for 10 min at ambient temperature. Plasma was separated from red blood cells and then subjected to an additional centrifugation at 18,000 g for 10 min at ambient temperature. Cell-free plasma was aliquoted and stored at −80°C. Extraction of cfDNA was performed using a fully automated QIAGEN platform, QIAsymphony SP, and QIAsymphony DSP Virus/Pathogen Midi Kit (QIAGEN). This is a bead-based custom protocol, optimized to work with 3 ml of plasma as starting material. The extraction process includes lysis, binding, wash, and elution steps. The final product was a 60 μl elution of cfDNA with an average size ~170-200 bp. Quality and quantity of cfDNA were evaluated with automated electrophoresis using a fragment analyzer with high sensitivity genomic DNA analysis kit (Advanced Analytical). A mean 122 ng of cfDNA was isolated at each timepoint and analyzed for wildtype and mutant IDH1 and IDH2 alleles by droplet digital PCR.

### Droplet digital PCR

An assay specific for each wildtype and mutant allele was designed and ordered through Biorad (IDH1 R132C: dHsaMDV2010053; IDH2 R172V: dHsaMDS874523342). Cycling conditions were tested to ensure optimal annealing/extension temperature as well as optimal separation of positive from empty droplets. All reactions were performed on a QX200 ddPCR System (Bio-Rad) and evaluated in technical duplicates. Reactions were partitioned into a median of ~16,000 droplets per well using the QX200 droplet generator and run on a 96-well thermal cycler. Plates were then analyzed with the QuantaSoft v1.7 to assess the number of droplets positive for mutant or wildtype DNA.

### Cell culture and DNA constructs

The chondrosarcoma cell line JJ012^25^ with an endogenous IDH1 R132C mutation was maintained at low passage number in high glucose DMEM with 10% FBS, glucose 25 mM, and glutamine 4 mM and split every 2-3 days before reaching confluence. Cultured cells repeatedly tested negative for mycoplasma throughout the experimental period. The IDH2 wildtype and R172V constructs were cloned by standard site-directed mutagenesis (Thermo) and Gibson Assembly (New England Biolabs) into pCDNA3.1 (Thermo) and verified by Sanger sequencing. For transient transfection experiments, cells were transfected using polyethylenimine. Two days after transfection, cells were treated with DMSO, ivosidenib (50 nM), or ivosidenib (50 nM) plus enasidenib (50 nM) and then harvested for protein and metabolite analysis one day later.

### Gel electrophoresis and western blotting

For denatured gel electrophoresis, cells were harvested in 1x RIPA buffer (Cell Signaling), centrifuged at 21,000 g at 4° C, and supernatants were collected. Cleared cell lysates were quantified by BCA assay (ThermoFisher) and normalized for total protein concentration. Samples were separated by SDS-PAGE, transferred to nitrocellulose membranes (Life Technologies), blocked in 5% milk prepared in Tris buffered saline with 0.1% Tween 20 (TBST), incubated with primary antibodies overnight at 4° C then horseradish peroxidase (HRP)-conjugated secondary antibodies (GE Healthcare; anti-mouse, NA931V, sheep, 1:5000; antirabbit, NA934V, donkey, 1:5000). After incubation with ECL (ThermoFisher or GE Healthcare), imaging was performed using the Amersham Imager 600 (GE Healthcare). Primary antibodies used included: anti-IDH1 (Proteintech, 12332-1-AP; rabbit; 1:1000), anti-IDH2 (Abcam, ab55271; mouse; 1:1000), and anti-vinculin (Sigma, V4505; mouse; 1:1000).

### Metabolite extraction and analysis

Metabolites were extracted from plasma (100 μl) or cell cultures (6-well plates) with ice-cold 80:20 methanol:water containing 2 μM deuterated 2-hydroxyglutarate (D-2-hydroxyglutaric-2,3,3,4,4-d_5_ acid; deuterated-2HG) as an internal standard. After overnight incubation at −80° C, plasma or cell extracts were sonicated and centrifuged at 21,000 g for 20 min at 4° C to precipitate protein. Extracts were then dried in an evaporator (Genevac EZ-2 Elite). Metabolites were resuspended by addition of 50 μl of methoxyamine hydrochloride (40 mg/ml in pyridine) and incubated at 30° C for 90 min with agitation. Metabolites were further derivatized by addition of 80 μl of N-methyl-N-(trimethylsilyl) trifluoroacetamide (MSTFA) + 1% 2,2,2-trifluoro-N-methyl-N-(trimethylsilyl)-acetamide, chlorotrimethylsilane (TCMS; Thermo Scientific) and 70 μl of ethyl acetate (Sigma) and incubated at 37° C for 30 min. Samples were diluted 1:2 with 200 μl of ethyl acetate, then analyzed using an Agilent 7890A GC coupled to Agilent 5975C mass selective detector. The GC was operated in splitless mode with constant helium carrier gas flow of 1 ml/min and with a HP-5MS column (Agilent Technologies). The injection volume was 1 μl and the GC oven temperature was ramped from 60° C to 290° C over 25 min. Peaks representing compounds of interest were extracted and integrated using MassHunter vB.08.00 (Agilent Technologies) and then normalized to both the internal standard (deuterated-2HG) peak area and plasma volume or protein content as applicable. Ions used for quantification of metabolite levels were 2HG *m/z* 247 (confirmatory ion *m/z* 349) and deuterated-2HG *m/z* 252 (confirmatory ion *m/z* 354). Peaks were manually inspected and verified relative to known spectra for each metabolite. Absolute metabolite quantitation was performed using an external calibration curve with deuterated-2HG internal standard and the resulting concentrations corrected for the total plasma volume extracted. For the patient with IDH1-mutant intrahepatic cholangiocarcinoma, plasma and tumor 2HG was measured using a qualified liquid chromatography-tandem mass spectrometry (LC-MS/MS) method with a lower limit of quantitation of 30 ng/ml for plasma or 100 ng/g for tumor as previously described^7^.

---

**Supplemental Figure 1.**
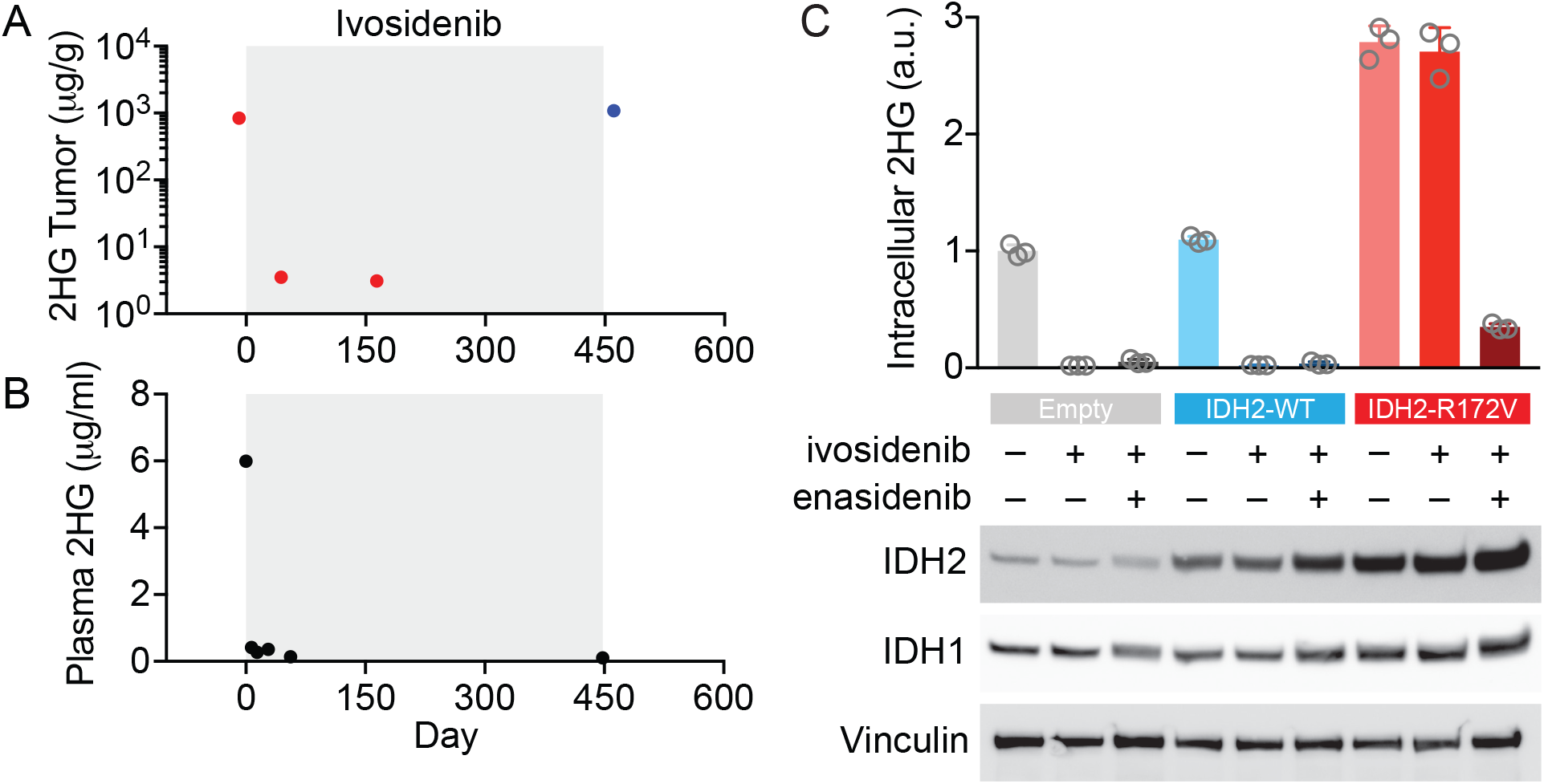
2HG levels in primary specimens from Case 3 and a cell line expressing both mutant IDH1 and mutant IDH2. (A) Tumor and (B) plasma 2-hydroxyglutarate (2HG) levels as measured by liquid chromatography–mass spectrometry (LC-MS) for the patient with IDH1 R132C-mutant intrahepatic cholangiocarcinoma (Case 3) treated with the mutant IDH1 inhibitor ivosidenib (gray box). For (A), red dots indicate biopsies from the primary tumor and the blue dot indicates a biopsy from the escape lesion as depicted in Figure 2. (C) An IDH1 R132C-mutant chondrosarcoma cell line (JJ012) was transfected with empty vector (Empty), IDH2 wildtype (IDH2-WT), or IDH2-R172V. Cells were treated for 24 hr with vehicle, ivosidenib (50 nM), or ivosidenib (50 nM) plus enasidenib (50 nM), and intracellular 2HG levels were measured by GC-MS. Error bars depict mean ±SEM for triplicate cultures. Western blot showing IDH2 and IDH1 protein levels. Vinculin serves as a loading control. Results for C are representative of 2 independent experiments.

**Supplemental Figure 2.**
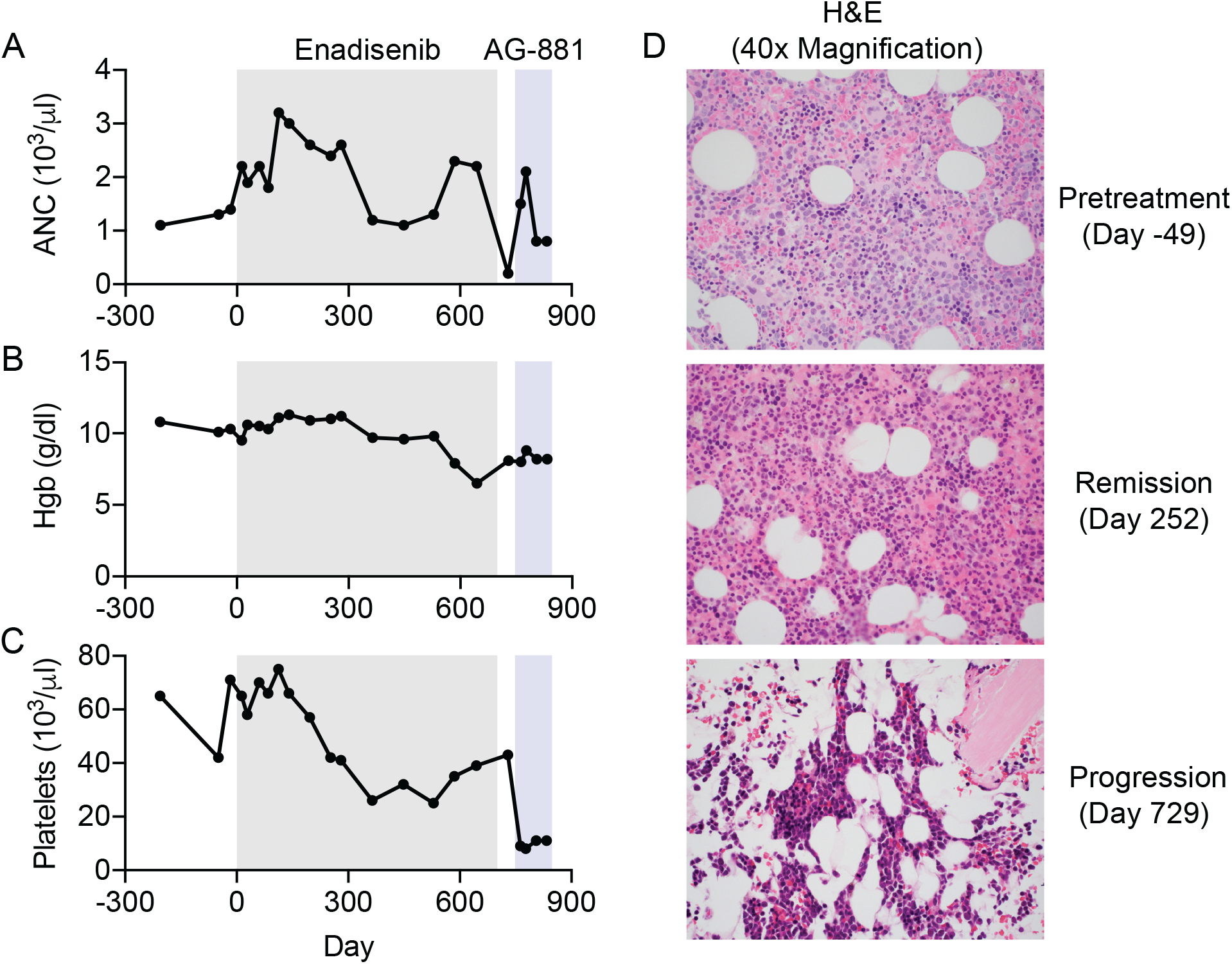
Additional clinical and laboratory features for the patient described in Case 4. (A) Absolute neutrophil count (ANC), (B) hemoglobin (Hgb) concentration, (C) platelet count, and (D) hematoxylin and eosin (H&E) staining of bone marrow at indicated timepoints in relation to treatment with the mutant IDH2 inhibitor enasidenib (gray box) and the dual IDH1/2 inhibitor AG-881 (blue box). Images show 40x magnification and are representative fields of a single bone marrow core biopsy performed at each time point.

